# A short ERAP2 that binds IRAP is expressed in macrophages independently from gene variation

**DOI:** 10.1101/2022.03.07.483178

**Authors:** Benedetta Mattorre, Silvana Caristi, Simona Donato, Emilia Volpe, Marika Faiella, Alessandro Paiardini, Rosa Sorrentino, Fabiana Paladini

## Abstract

The M1 zinc metalloproteases ERAP1, ERAP2 and IRAP play a role in HLA-I antigen presentation by refining the peptidome either in the ER (ERAP1 and ERAP2) or in the endosomes (IRAP). They have been also entrusted with other, although less defined, functions such as the regulation of the angiotensin system and blood pressure. In humans, ERAP1 and IRAP are commonly expressed. ERAP2 instead has evolved under balancing selection that maintains two haplotypes one of which undergoing RNA splicing leading to nonsense-mediated decay and loss of protein. Hence, likewise in rodents in which the ERAP2 gene is missing, about a quarter of the human population does not express ERAP2. We report here that macrophages, but not monocytes or other mononuclear blood cells, express and secrete an ERAP2 shorter form independently from the haplotype. The generation of this “short” ERAP2 is due to an autocatalytic cleavage within a distinctive structural motif and requires an acidic microenvironment. Remarkably, ERAP2 “short” binds IRAP and the two molecules are co-expressed in the endosomes as well as in the cell membrane. Of note, the same phenomenon could be observed in some cancer cells. These data prompt to reconsider the role of ERAP2 which might have been maintained in humans because fulfilling a relevant function as “short” form in specialized cells.

## INTRODUCTION

The aminopeptidases ERAP1, ERAP2 and IRAP, the latter being the product of the gene *LNPEP*, are members of the oxytocinase subfamily of M1 Zinc-metallopeptidases whose corresponding genes lie contiguously on chromosome 5 (1,2). ERAP1 and ERAP2 reside in the ER where they co-operate to trim the N-terminal peptide residues to the correct length to bind the HLA-I molecules (3). IRAP instead, thanks to an additional N-terminal cytoplasmic domain, is retained in the endosomal vesicles from where it can traffic to the cell membrane forming a type II integral membrane glycoprotein (4). IRAP is a multifaceted protein: it has been shown to be involved in cross-presentation in the endosomes of dendritic cells (DC) and, when in the cell membrane, to catalyze the final step of the angiotensinogen to angiotensin IV (AT4) conversion being itself a receptor for AT4 (5,6). In addition, it has been shown to be involved in several other functions, ranging from the insulin metabolic pathway to vesicular trafficking and even in cognitive processes (7,8). Furthermore, the three genes have been found associated with autoimmune and inflammatory conditions, hypertension, and cancer (2, 9-15). Since they are regulators of the Renin-Angiotensin system (RAS), their imbalance can have indeed consequences on different aspects of the associated pathologies (16-18). In the MHC-I-opathies (19), while *ERAP1* association resides in variants influencing its trimming activity, the molecular terms of the association of *ERAP2* and *LNPEP* with some of these diseases are at present poorly understood (20). In particular, *ERAP2* has been shown to be involved in MHC-I-opathies and other conditions, while much of its biology remains uncertain, including the dramatic variability in its functional opposite haplotypes maintained in the population. This is mainly due to a balanced polymorphism at SNP rs2248374 that makes one haplotype null due to a shift in the splicing that leads to mRNA instability and non-sense mediated decay (NMD). Consequently, about a quarter of the human population does not express ERAP2 full length and about 50% shows an allelic exclusion. Of note, this gene is missing in rodents as well as in several other species (2, 21). These observations raise several questions about the physiological role of ERAP2. Indeed, by looking at the expression of ERAP1, ERAP2 and IRAP along the zoological scale, we have recently pointed out how a high degree of redundancy and, most likely interchangeability, characterizes these three aminopeptidases (2). ERAP1 has been shown to be secreted (22) and IRAP has also a soluble counterpart detectable in the serum of pregnant women (23). More recently, an ERAP2 soluble isoform has been described as released by human monocyte-derived macrophages (MDMs) in response to IFNγ/LPS stimulation and this corresponds to an increased CD8+ T cells activity (24). With the aim to gain additional information about the interconnections among the three aminopeptidases, we have investigated their expression and interactions in different contexts. Remarkably, we observed an ERAP2 shorter subunit expressed by macrophages as well as by some cancer cells and which forms a complex with IRAP. This ERAP2 “short” corresponds to a N-terminal fragment that requires an acidic microenvironment to be generated and it is present even in those subjects expected to be ERAP2-deficient. These suggestive findings prompted us to reconsider the role of ERAP2 in physiology as well as in pathology.

## MATERIALS AND METHODS

### Cell Lines

Human histiocytic leukemia U937 (ATCC: CRL-1593.2, Manassas, VA, USA) cells were maintained in culture (10^6^ cells/ml) in Roswell Park Memorial Institute medium (RPMI 1640, Invitrogen) containing 10% of heat inactivated fetal bovine serum (FBS, Invitrogen) and supplemented with 10 mM Hepes (Gibco, #15630-056), 1 mM pyruvate (Gibco, #11360-039), 2.5 g/l D-glucose (Merck). U937 cells were differentiated to macrophages by 24 h incubation with 20 nM phorbol 12-myristate 13-acetate (PMA, Sigma, P8139) followed by 24 h incubation in RPMI medium. Macrophages were polarized to M1 by incubation with 20 ng/ml of IFN-γ (R&D system, #285-IF) and 10 pg/ml of LPS (Sigma, #2630). Macrophage M2 polarization was obtained by incubation with 20 ng/ml of IL-4 (R&D Systems, #204-IL) and 20 ng/ml of IL-13 (R&D Systems, #213-ILB). U937 cells were plated in complete RPMI 1640 for each condition. M2-like U937 phenotype was obtained also by treating the cells with 80 nM PMA for five days.

Caco-2 (ATCC: HTB-37), LoVo (ATCC: CCL-229), HEK293T (ATCC: CRL-1573) and HeLa (ATCC: CCL-2) were cultured in completed Dulbecco’s Modified Eagle Medium (DMEM, Invitrogen) supplemented with 10% FBS; K-562 (ATCC CRL-3343), C1R (ATCC: CRL-2369) and HL-60 (ATCC: CRL-2257) were grown (10^6^ cells/ml) in RPMI 1640 supplemented with 10% FBS.

### Antibodies

The following mouse monoclonal antibodies were used: anti-ERAP1 (mAb clone B-10, sc-271823 Santa Cruz), anti-ERAP2 (mAb clone 3F5, MAB 3830 R&D Systems) (25), anti-IRAP (mAb F-5 sc-365300, Santa Cruz), anti-β-Actin (mAb clone C4, sc-477778 Santa Cruz), anti-CD14 (mAb UCHM1, Abcam); anti-MS4A4A (mAb MAB7797-SP, R&D); anti-EEA1 (mAb MA5-31575, Invitrogen).

### Macrophages-Derivated-Monocytes (MDMs) generation and CD14 negative PBMCs

Human Peripheral Blood Mononuclear Cells (PBMCs) were purified from peripheral blood of anonymous donors from the local data banks by density centrifugation on Lympholyte (Cedarlane Laboratories). The study was carried out in accordance with the recommendation of Ethical Committee of the Policlinico Umberto I (Sapienza University, Rome, Italy). All subjects gave written informed consent in accordance with the Declaration of Helsinki (ethical code N. 1061bis/2019, 13/09/2019).

Monocyte-and lymphocyte-enriched PBMC suspensions were sorted using Isolation kits and Depletion Column Type LS (Miltenyi Biotec Inc., Auburn, CA) according to the supplier’s instructions. Cell viability, as measured by Trypan blue exclusion, always exceeded 95%. Monocytes and lymphocytes isolated by this procedure were found by FACScan flow cytometer (FACsort, Becton-Dickinson) analysis more than 90% and 98% pure respectively. Cells were adjusted to the density of 10^6^ cells/ml and were suspended in RPMI 1640 supplemented with 10% heat-inactivated FBS, 2mM L-glutamine, 25 U/ml penicillin and 25 U/ml streptomycin (all purchased from GIBCO) in cell culture plates and kept at 37°C in a humidified 7% CO_2_ incubator.

For MDMs differentiation, purified cells were cultured in complete medium added with 100 ng/ml macrophage colony-stimulating factor (M-CSF) (Peprotech, Stockholm, Sweden) or 100 ng/ml granulocyte–macrophage colony-stimulating factor (GM-CSF) (Peprotech), for 6 days. Media were changed every 2–3 days. After seven days, MDMs were stimulated with either LPS (100 ng/ml) and IFNγ (20 ng/ml) (M1-stimulation) or IL-4 (20 ng/ml) (M2-stimulation) or left untreated (M0) for 24 hours before collecting the cells and the respective surnatants.

CD14 negative PBMCs were cultured at 5 × 10^6^ cells/ml in RPMI 1640 with 10% FBS in the absence or presence of Phytohemagglutinin (PHA) (1 μg/ml) (Roche) for 24 h.

Cell and bovine serum albumin-free surnatants were harvested by centrifugation (1000 g; 15 min) and stored at 4°C until assayed.

### DNA extraction and rs2248374 genotyping

Genomic DNA from EDTA-treated peripheral blood samples were extracted using QIAamp DNA Blood mini-kit (Qiagen, Hilden, Germany) according to the manufacturer’s protocol. Genotyping of the SNP rs2248374 was performed by quantitative Real-Time Polymerase Chain reaction (qRT-PCR) with functionally tested TaqMan Allelic Discrimination Assay (C_25649529_10; 7300 real-time PCR system, Applied Biosystems).

### Flow cytometry analysis

Collected cells were washed and resuspended in staining buffer (PBS 1X +1% BSA) with Fc Receptor Binding Inhibitor Antibody (Invitrogen, # 14-9161-73) for 15 minutes at 4°C. After blocking, samples were incubated in the dark for 1 hour at 4 °C with saturating concentration of anti-human primary mouse monoclonal antibodies. After two washes in PBS 1X +1% BSA, cells were incubated with secondary antibodies: Alexa Fluor 488® F(ab’)2 fragment of goat anti-mouse IgG (H+L) (Invitrogen) and/or Alexa Fluor 594® F(ab’)2 fragment of goat anti-mouse IgG (H+L) (Invitrogen). Samples were then washed with PBS 1X+1% BSA and resuspended in 200 μl of the same buffer. Macrophages were electronically gated according to light scatter properties to exclude cell debris and contaminating lymphocytes. Fluorescence was measured using a FACSCalibur flow cytometer (Becton Dickinson, Missisauga, ON, Canada) and analyzed using FlowJo software (Tree Star Inc., Ashland, OR, USA).

### Western Blot

Approximately 5 × 10^6^ cells for conditions were harvested, washed twice in cold 1x PBS and resuspended in 100 μl of hypotonic buffer solution (20 mM Tris-HCl, pH 7.4; 10 mM NaCl; 3 mM MgCl_2_) containing 100 U/ml of phenylmethylsulfonyl fluoride (PMSF), 1 μg/ml of aprotinin, 0.5% sodium deoxycholate, proteinase inhibitors cocktail (Pierce) and 25 μl of 10% NP40. After centrifugation at 16000 *g* for 15 min at 4 °C, total protein concentration was determined by the Biorad protein assay kit (Biorad, Hercules, CA) with BSA used as standard. Forty μg of protein extract for each sample were separated on a 4–12% NuPage Bis-Tris gel (Invitrogen) at 125 V for 100 min in NuPage MES SDS Running Buffer (Invitrogen)and transferred to nitrocellulose membranes.

For each sample, 200 µl of BSA-free surnatant was recovered and proteins were precipitated by chloroform/methanol procedure (26). The proteins were resuspended in denaturing loading Buffer 2X, boiled for 5 min, separated on a 4–12% NuPage Bis-Tris gel (Invitrogen) and transferred to nitrocellulose membranes. After incubation ON with the respective antibodies, the membranes were washed twice in 0.1% Tween PBS, incubated 1h with horseradish peroxidase–conjugated secondary Ab (Jackson Immunoresearch Laboratories, Inc. West Grove, PA) and revealed by ECL Western blotting detection system (Amersham). The proteins were visualized by ChemiDoc XRS+ System (Biorad, California, USA) (maximum exposure: 600 seconds).

### Confocal microscopy

Cells were washed with PBS 1X + BSA 1%, incubated with Fc Receptor Binding Inhibitor Antibody for 15 minutes at 4°C, fixed and permeabilized with EtOH (70%) or with paraformaldehyde (4%) for 30 minutes. Any specific bond was saturated by incubating the samples with PBS+BSA 1% for 30 minutes. After two washes, cells were incubated with primary antibodies for 45 minutes and then with secondary antibodies (Alexa Fluor 594® F(ab’)2 fragment of goat anti-mouse IgG (H+L) and Alexa Fluor 488® F(ab’)2 fragment of goat anti-mouse IgG (H+L), Invitrogen) for 45 min. Laser lines: LASER ARGON 488 for Alexa 488 (green), LASER A HELIUM (He) 543 for Alexa 594 (red). The images were processed with the ImageJ software (27). Colocalization was determined based on the overlap coefficient according to Manders automatically calculated by ZEISS Zen Software. The overlap coefficient has a range from +1 (perfect correlation) to 0 denoting no relationship. Values >0.5 represent the high probability that there is an overlap denoting colocalization, while values closer to 0 (<0.5) denote a lower probability that pixels from both channels, in relation to the entire image, have overlapped (no relationship) (28).

### Immunoprecipitation

Immunoprecipitation was carried out according to the PierceTM Crosslink protocol Immunoprecipitation kit (Thermo Scientific). In short, the antibodies were “crosslinked” to the resin using the DSS (disuccinimidyl suberate) 0.25mM reagent in DMSO (dimethyl sulfoxide). For each immunoprecipitation, 1mg of total protein lysate was previously incubated with control resin, in order to obtain an INPUT free of non-specific bonds. The INPUT was then incubated with the antibody-bound resin ON at 4°C and the bound proteins eluted using the elution buffer kit.

### Transfection

HEK293T cells were seeded in 12-well plates in complete medium (DMEM) containing 10% FBS and transfected after 24 hours with jetPEI® (Polyplus-transfection ® SA) according to the manufacturer’s protocol. A total of 1 μg plasmid DNA/well was transfected: empty vector pcDNA 3.1 or plasmid constructs carrying the first 1605bp nucleotide sequence of the ERAP2 “G” haplotype or the “A” haplotype encoding the first 532 or 534 amino acids respectively (Figure S1). The difference in lenght is due to the presence of a stop codon in the “G” haplotype. The plasmid construct was supplied by GenScript USA Inc.

### Modelling

The crystal structures of the soluble domains of human ERAP1 (PDB 2YD0) (29), ERAP2 (PDB 5AB0) (30) and IRAP (PDB 4PJ6) (31) were used for structural analysis and comparison. The HADDOCK server with default parameters was used for ∼55 kDa ERAP2/IRAP docking (32). The Prodigy algorithm was used to predict the binding affinity and for the analysis of the interface (33,34).

Rosetta 3.13 Linux version was used to predict the conformation of ERAP2 short isoforms in the presence of the LAFLGENA and the VRIKRVTE motifs. The initial atomic coordinates were taken from the X-ray crystal structure of ERAP2 (Protein Data Bank ID: 5AB0). The Python protocol described in Yang et al. (35), which makes use of the Rosetta 3.13 neural network to generate inter-residue distance and orientation constraints for a sequence of unknown structure, was used. The Rosetta minimizer was then used to find the backbone conformation consistent with the constraints.

### Quantification and statistical analysis

ImageJ, ChemiDoc XRS+ System and FlowJo software were used to for image analysis.

## RESULTS

### Monocyte-derived Macrophages express a short ERAP2 isoform that binds IRAP

Peripheral blood mononuclear cells (PBMC) from six donors, three genotyped as G/G and three as A/A at the allelic variant rs2248374, were analyzed for the expression of ERAP1, ERAP2 and IRAP. The CD14+ positive mononuclear cells were then differentiated to either M1 or M2 macrophages whereas the corresponding CD14 negative mononuclear cells were activated by phytohaemagglutinin (PHA). To obtain type 1 and type 2 macrophages, monocytes were kept in culture for seven days with either GM-CSF or M-CSF, and then stimulated for 24 hours with either IFNγ and LPS (M1) or IL-4 (M2). The expression of the three aminopeptidases was analyzed by western blot (Figure 1A and 1B). ERAP2, as expected, was expressed by those subjects genotyped as A/A but not by those genotyped as G/G at SNP rs2248374. Interestingly however, the blotting with the anti-ERAP2 MoAb revealed the presence of a protein of ∼55 kDa in the differentiated monocytes, independently from the stimulus and from the SNP rs2248374 genotyping. Of note a band corresponding to the ∼55kDa was also observed in the surnatants of the MDMs, in particular in the case of M2-type. Since M2 macrophages are characterized by the expression of MS4A4A (36), this marker was used to analyze the MDMs population for its expression. In our hands, the freshly isolated monocytes from different individuals showed a variable percentage of MS4A4A positive cells (Figure S2). This can be due to different reasons: their native state, the stimuli received during the isolation or through the CD14 positive selection etc. However, once differentiated, MS4A4A positivity raises up to 85-90% in the case of M2 and to about 70-80% in the case of M1. We therefore investigated whether the secreted ERAP2 ∼55kDa protein was also detectable in the cell surface of MDMs. The results showed that a high percentage of the MS4A4A positive MDMs stained also positive for ERAP2 (Figure 1C) whereas CD14 negative PBMC did not express appreciable levels of either MS4A4A or ERAP2 even after seven days of PHA-stimulation. The positivity for ERAP2 in the MDMs derived from the individuals genotyped as G/G at rs2248374, indicates that it is the “short” and not the full length ERAP2 that is present in the cell membrane. In addition, the co-expression of the MS4A4A and ERAP2 strongly suggests that the “short” ERAP2 is secreted by the M2-type macrophages since, in our hands, MS4A4A-positive cells are present also in the putative M1-type.

**Figure 1.**
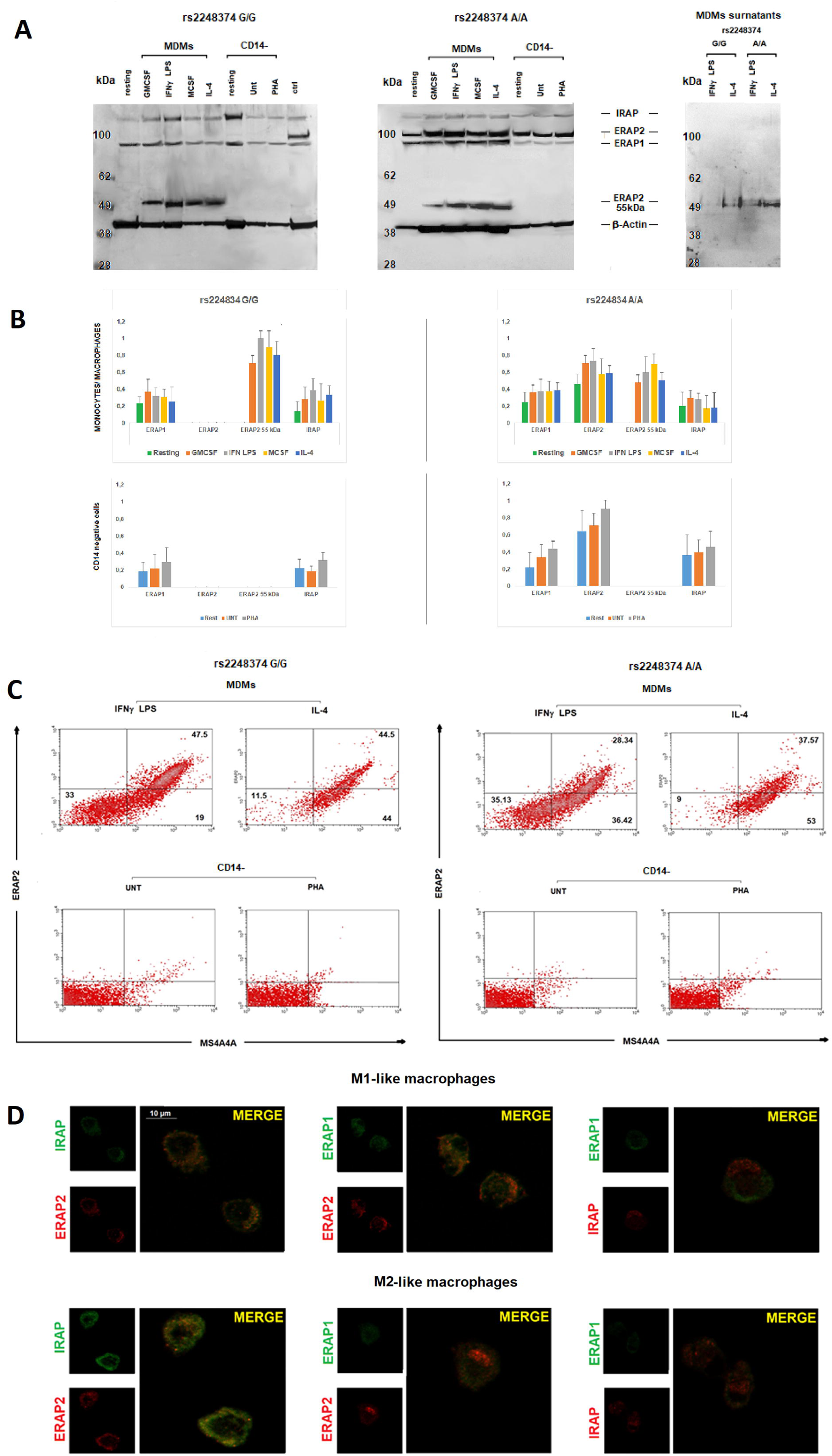
Monocyte-Derived-Macrophages express a ∼55kDa ERAP2 form. **(A)** Western Blot analysis of ERAP1, ERAP2 and IRAP expression in monocytes, M1 and M2 macrophages and CD14 neg PBMCs, the latter treated or not with PHA, from two donors genotyped as G/G (left) or A/A (middle) at rs2248374. Right: surnatants from MDMs treated with either IFNγ+LPS (M1) or IL-4 (M2). The images are representative of six experiments using PBMCs derived from three donors genotyped as G/G and three as A/A at rs224837. Control (Ctrl): protein extract from B-LCL (EBV^+^ B-lymphoblastoid cell line genotyped A/A at rs2248374). **(B)** Monocytes/macrophages and CD14 negative PBMC western blot densitometric analysis: the intensity of bands was quantified using the program UN-SCAN-IT gel (Silk Scientific Inc., Orem, UT). Results were evaluated as ratio of intensity between relevant protein and β Actin from the same sample. Values are the mean+ SD of three independent experiments. **(C)** Cytofluorimetric analysis of the same samples as in **A**. M1 (IFNγ+LPS treated) and M2 (IL-4 treated) macrophages were analyzed for the expression in the cell membrane of ERAP2 and of the M2 marker MS4A4A. The image is representative of three different experiments. **(D)** Confocal microscopy images of M1 (IFNγ+LPS treated) or M2 (IL-4 treated) MDMs generated from monocytes from a subject genotyped as G/G at rs2248374. Cells were fixed, permeabilized and co-stained with a two-by-two combinations of ERAP1, ERAP2 and IRAP antibodies. Left: ERAP2 (red) and IRAP (green) Middle: ERAP2 (red) and ERAP1 (green) right: ERAP1 (green) and IRAP (red). Images were obtained using the 63× objective with a 3x magnification (z=3-4μm). Co-localization was determined based on the overlap coefficient according to Manders automatically calculated by ZEISS Zen Software Scale bar:10 μm. The image is representative of three different experiments.

The confocal analysis of the three aminopeptidases on MDMs (Figure 1D) revealed that, of all the possible two by two combinations of the three aminopeptidases, a co-localization of ERAP2 with IRAP was evident, particularly in the case of M2-type macrophages (overlap coefficient = 0.73 vs M1 overlap coefficient = 0.56) Of note, MDMs shown here derive from a donor genotyped as G/G at rs2248374 and therefore not expressing the full length ERAP2. These observations allowed us to conclude that ERAP2 “short” co-localizes with IRAP both inside the cell as well as in the cell membrane. No co-localization between IRAP and ERAP1 or between ERAP1 and ERAP2 could be observed (overlap coefficient < 0.50).

These data indicate that, independently from the *ERAP2* genotype, MDMs express a “short” ERAP2 that co-localizes with IRAP and that it is secreted primarily by the M2-type macrophages.

### Differentiation of U937 cells correlates with the loss of ERAP2 full length and the occurrence of a ∼55kDa fragment

To get more insights into these unexpected findings, we used the human pro-monocytic myeloid leukemia cell line U937 as a model for macrophage differentiation. After 48h of treatment with Phorbol 12-myristate (PMA), the adherent cells underwent either LPS plus IFN-γ, or to IL-4 plus IL-13 treatment to induce an M1-like or an M2-like polarization, respectively. The expression of ERAP1, ERAP2 and IRAP was therefore analyzed by immunofluorescence (Figure S3) and by western blotting (Figure 2A). Compared to the untreated cells, PMA-treated cells show a higher expression of MS4A4A and, in parallel, of ERAP2 whereas ERAP1 and IRAP did not vary (Figure S3). Western blot analysis (Figures 2A and 2B) showed that U937 cells, despite being genotyped as G/G at SNP rs2248374, express the canonical ERAP2 protein (115 kDa). This was not surprising since it has been already reported in other transformed cell lines carrying the G/G genotype (37), some of which showed here (Figure S4), indicating that the alternative splicing associated with the presence of the variant G at rs2248374 is not a universal occurrence. PMA treatment *per se* led to a substantial decrease in the expression of the ERAP2 canonical protein replaced by a fragment of ∼55 kDa. An additional band of ∼65 kDa is expressed by the U937 as well by other cancer cell lines (Figure S4) which however was not observed in MDMs and therefore was not further investigated here. Additional differentiation of the U937 cells by M1 or M2 stimuli did not substantially modify the expression of the different isoforms. Likewise to MDMs (Figure 1), the protein of ∼55 kDa was also detected in the surnatant, in particular from M2-like or PMA-treated cells. ERAP1 was expressed across all conditions and, as previously described, it was secreted as full length upon stimulation (22). IRAP instead, could not be detected in the surnatant, either because not secreted by these cells or because the MoAb (F-5 sc-365300, Santa Cruz) recognizes a region proximal to the cell membrane (data not shown).

**Figure 2.**
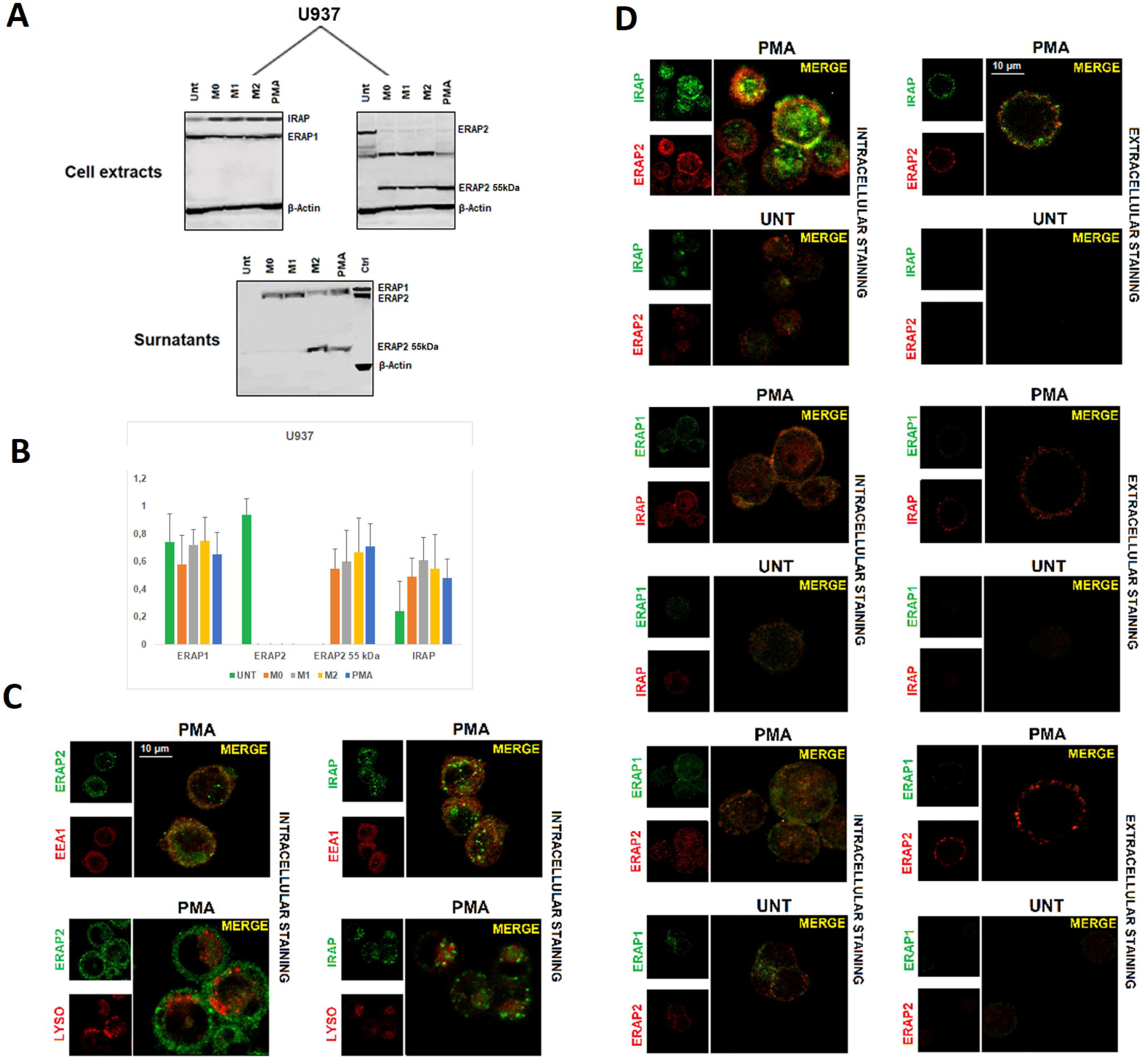
ERAP1, ERAP2 and IRAP expression in U937 cell line. **(A)** Western Blot analysis of ERAP1, ERAP2 and IRAP expression in U937 cells untreated or differentiated to macrophages by different stimuli (see text): membranes were blotted with anti-ERAP1, anti-ERAP2 and anti-IRAP MoAbs. Images of anti-ERAP2 are shown separately to highlight the bands specific for the anti-ERAP2 MoAb. The respective surnatants were blotted with the three antibodies (IRAP not shown). Control (Ctrl): protein extract from B-LCL (EBV^+^ B-lymphoblastoid cell line genotyped A/A at rs2248374). Results shown are representative of 3 independent experiments. **(B)** Densitometric analysis: the intensity of bands was quantified using the program UN-SCAN-IT gel (Silk Scientific Inc., Orem, UT). Results were evaluated as ratio of intensity between relevant protein and β Actin from the same sample. Values are the mean+ SD of three independent experiments. **(C)** Confocal images of two-by-two combinations of ERAP1, ERAP2 and IRAP in U937 cells treated or not with PMA. Colocalization was determined based on the overlap coefficient according to Manders automatically calculated by ZEISS Zen Software. Left panel: intracellular staining of permeabilized U937 cells. Right panel: extracellular staining of permeabilized cells. Results shown are representative of 3 independent experiments. **(D)** Images show the co-localization of ERAP2 with IRAP either in the endosomes (upper panel) or in the lysosomes (lower panel). The markers used are EEA1 for the endosomes and LysoTracker® Red DND-99-positive for the lysosomes.

PMA-differentiated U937 cells alongside with an untreated control were kept in culture for 5 days and then analyzed using confocal microscopy. Indeed, following PMA-treatment, these cells acquire a M2-like phenotype as revealed by the expression of the MS4A4A marker (Figure S3) (36,38). We therefore proceeded to analyze the localization of ERAP1, ERAP2 and IRAP. The data showed an intracellular co-localization of ERAP2 with IRAP (overlap coefficient = 0.79) but not with ERAP1 (overlap coefficient < 0.5) (Figure 2B). As in the case of MDMs, no co-localization of IRAP with ERAP1 or of ERAP1 with ERAP2 was appreciable (overlap coefficient <0.5). The staining with the endosomal marker EEA1 showed that the colocalization takes place specifically in the endosomes where IRAP1 resides (Figure 2C) (overlap coefficient =0.74). Interestingly, IRAP and ERAP2 colocalize in the cell membrane as well (overlap coefficient =0.67). Given that the differentiated U937 cells do not express the full length ERAP2 protein (Figure 2A), we could confirm that the co-localization with IRAP specifically refers to the ERAP2 ∼55kDa. To verify whether this co-localization was a special feature of the MDMs or if the presence of the “short” ERAP2 was a sufficient condition, we took advantage of two colon carcinoma cell lines, Caco-2 and LoVo. Indeed, Caco-2 spontaneously express the ∼55kDa ERAP2 whereas LoVo cells do not (Figure S4). We therefore analyzed its expression. As show in Figure 3, Caco-2, but not LoVo cells, not only stained positive for ERAP2 in the cell surface (Figure 3A), but the confocal imaging showed, as in the case of MDMs, a co-localization of IRAP with ERAP2 in Caco-2 (overlap coefficient = 0.71) but not in LoVo cells (overlap coefficient < 0.5) (Figure 3B). No soluble ERAP2 could be detected in the surnatant of either cell line (not shown). Once again, these data confirm that the “short” ERAP2 colocalizes with IRAP.

**Figure 3.**
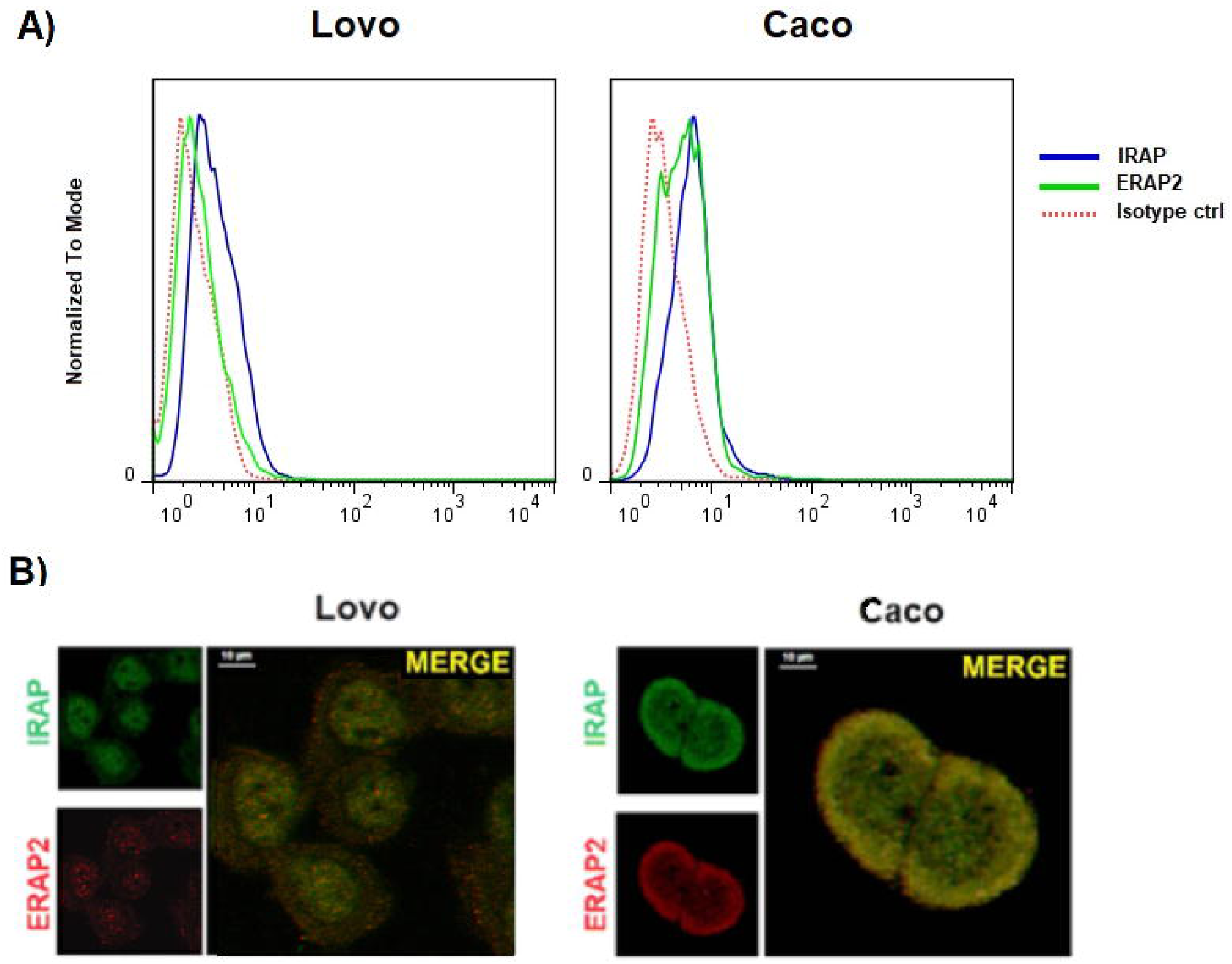
ERAP2 and IRAP expression in LoVo and Caco-2 cell lines. **(A)** Cytofluorimetric analysis of the cell membrane expression of ERAP2 (green) and IRAP (blue). LoVo cells express no ERAP2 in the cell membrane whereas CaCo cells do. IRAP is equally expressed in both cells. **(B)** Confocal analysis of the intracellular expression of ERAP2 (red) and IRAP (green) in the same samples. Scale bar:10 μm. Results shown are representative of 3 independent experiments.

### IRAP directly binds the ∼55kDa ERAP2

Next, we investigated the nature of the ERAP2 and IRAP association by performing co-immunoprecipitation experiments. Cellular extracts from untreated or PMA-treated U937 cells underwent immunoaffinity with anti-ERAP2, anti-IRAP and, as control, anti-ERAP1 MoAbs. The eluates were then separated on SDS-PAGE and blotted with the three antibodies (Figure 4). As expected, each MoAb immunoprecipitated the corresponding protein in the untreated cells. Interestingly, in the PMA-treated cells, a band recognized by the anti-IRAP MoAb could be observed in the outflow of the *α*-ERAP2 column (Figure 4A first lower panel, lane 2). The reciprocal was also true and a band of ∼55kDa was recognized by the anti-ERAP2 MoAb when the lysate was immunoprecipitated with the anti-IRAP MoAb (Figure 4A second lower panel, lane 3). No co-immunoprecipitation between ERAP1 and either ERAP2 or IRAP was detectable (Figure 4A). As control, in each box the three MoAbs used were run in the last three lanes to exclude that the heavy chain of the MoAbs used in the immunoprecipitation assay, whose MW is ∼50kDa, could overlap with the ∼55kDa ERAP2. Of note, to elute ERAP2 from the beads conjugated with the corresponding antibody, a harsher treatment was necessary. In this case, the same ∼55kDa fragment was present in the eluates of both untreated and PMA-treated U937 cells, although the untreated cells expressed none or a barely detectable ∼55kDa fragment (Figure 2A) suggesting that the treatment to which the U937 cell line underwent in the co-immunoprecipitation procedure (i.e. the low pH of the elution buffer) caused the generation of the ∼55kDa fragment. To verify this, U937-untreated cell extracts underwent a pH gradient (from 8 to 5) before loading (Figure 4B). The band corresponding to the ∼55kDa fragment, absent at pH 8 and pH 7, is detectable at the lower pHs inversely correlating with the full length band (Figure 4B). We concluded that the ∼55kDa is generated from the full-length protein through a mechanism that requires acidic conditions.

**Figure 4.**
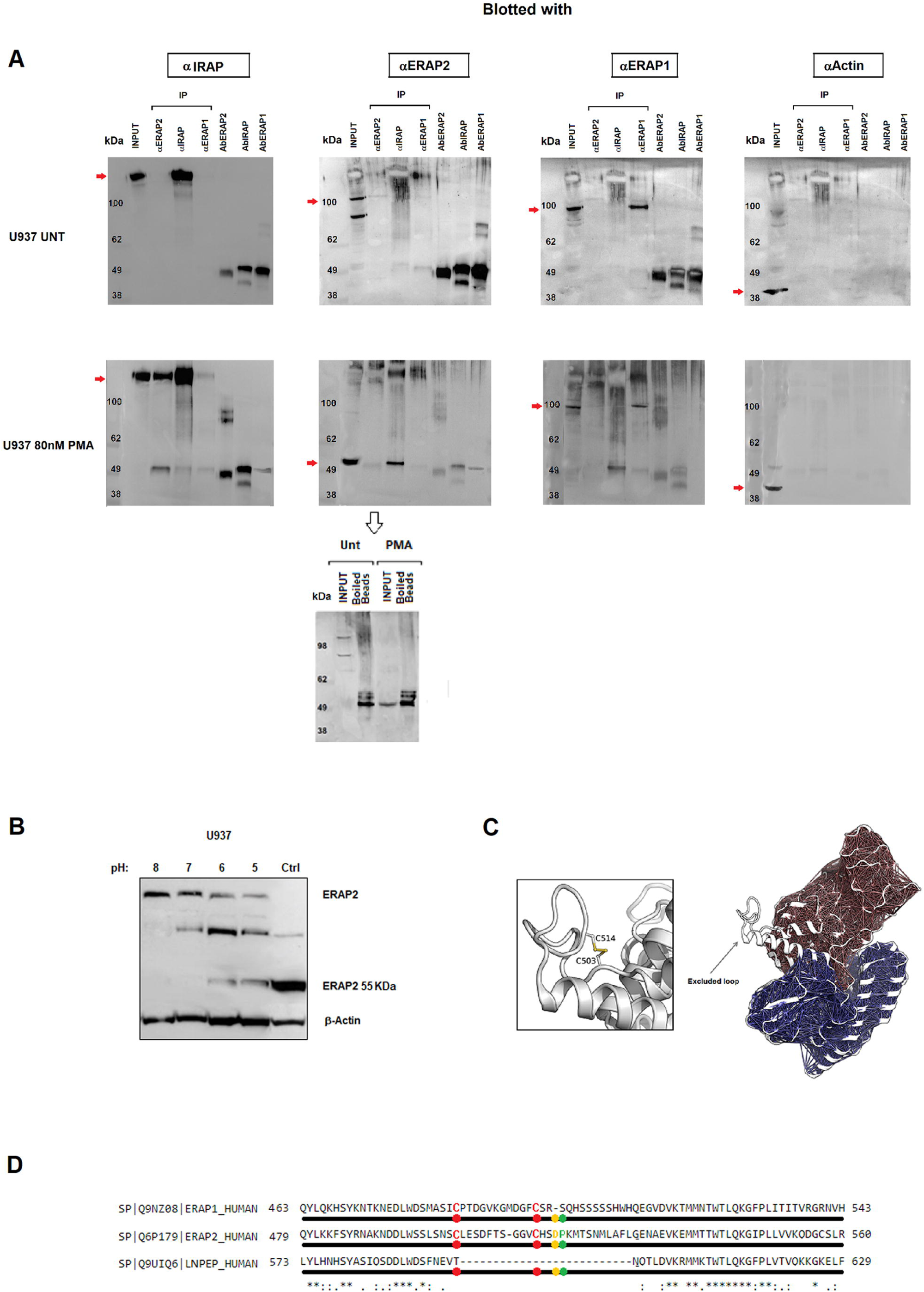
Immunoprecipitation of ERAP1, ERAP2 and IRAP. **(A)** Top membranes: Untreated U937 cells; bottom membranes: U937 treated with 80 nM PMA for 96h. Panels show the input and the immunoprecipitated with respectively anti-ERAP2, anti-IRAP, anti-ERAP1 and the respective antibodies run as controls in the last three lanes. From the left: the membranes were blotted with anti-IRAP, anti-ERAP2, anti-ERAP1 and anti-actin. In the PMA-treated series, the first panel has been blotted with the anti-IRAP which is present in the input (1° lane), in the immunoprecipitated with anti-IRAP (4° lane) but also in the immunoprecipitated with anti-ERAP2 (3° lane). The second panel has been blotted with the anti-ERAP2 antibody and the “short” ERAP2 is evident in the input and in the anti-IRAP immunoprecipitated. Of note, in the corresponding upper panel, accordingly to Figure 2, the input shows the two bands characteristic of the U937 cells whereas in the lower panel the input shows only the “short” ERAP2. The third panel shows the same samples blotted with the anti-ERAP1 antibody: ERAP1 is present in the input and in the anti-ERAP1 immunoprecipitated with no difference between the upper and lower panel. In the fourth panel, the immunoprecipitated samples have been blotted with an anti-actin as control. The last three lanes in each gel contain, as indicated, the three monoclonal antibodies used in the experiment. At the bottom: since the acid elution buffer did not allow the release of the ERAP2 bound to the beads (lane 2 of the second panel), they were heated (99°C, 5 min) to confirm the presence of the ERAP2 which was released as ∼55kDa fragment in both untreated and PMA-treated samples. **(B)** Untreated U937 cell lysates underwent a pH gradient and run on SDS-PAGE. CTRL=PMA-treated U937 cells **(C)** Structure of ERAP2 (PDB 5AB0) showing the extended loop that characterizes ERAP2 in comparison with ERAP1 (PDB 2YD0) and IRAP (PDB 4PJ6) (not shown). **(D)** Amino acid sequences of ERAP1, ERAP2 and IRAP in the region surrounding the cysteine bridge. The specific Asp-Pro motif site that characterizes ERAP2 is shown.

### ERAP2 contains a potential autocatalytic cleavage site

In order to gain structural insights into the mechanism of the cleavage observed in ERAP2 but not in the closely related ERAP1 and IRAP, the crystal structure of ERAP2 (PDB: 5AB0) was investigated and compared to the other two aminopeptidases (39). In the case of ERAP2, the protein structure shows the presence of a unique extended loop, encompassing residues 500-520 (Figure 4C). This distinctive structural motif is peculiar of ERAP2 and contains two potential sites, which could be sensitive to low pH regulation: 1) a disulfide bridge formed by Cys 503 and Cys 514, which might be broken at low pH values and 2) a specific Asp-Pro motif (Figure 4D). However, while the disulfide bridge is also conserved in ERAP1, the Asp-Pro motif is unique of ERAP2. This site has been reported to be particularly sensitive to pH variations (40) and it could represent a weak structural site that, in an acidic microenvironment, can determine the breakage thus generating the N-terminus ∼55kDa fragment recognized by the anti-ERAP2 MoAb.

A possible model of the ∼55kDa fragment of ERAP2 bound to IRAP was also investigated by protein-protein docking means, to tentatively explain the strong affinity observed in the co-immunoprecipitation experiments. As shown in Figure 5, ERAP2 is predicted to form a pseudo-symmetric heterodimer with IRAP, with the N-terminal β-sheet rich region of ERAP2 (residues 50-170) making extensive contacts with the corresponding region of IRAP. The predicted ΔG of binding and dissociation constant (Kd) values were -15.5 Kcal/mol and 4.3e^-12^, respectively, indicative of a strong affinity between the two proteins. Moreover, the large amount of surface area buried upon interaction and the complementarity of surface charge validate a plausible mechanism of dimerization, which could account for the specific interaction observed *in vitro*.

**Figure 5.**
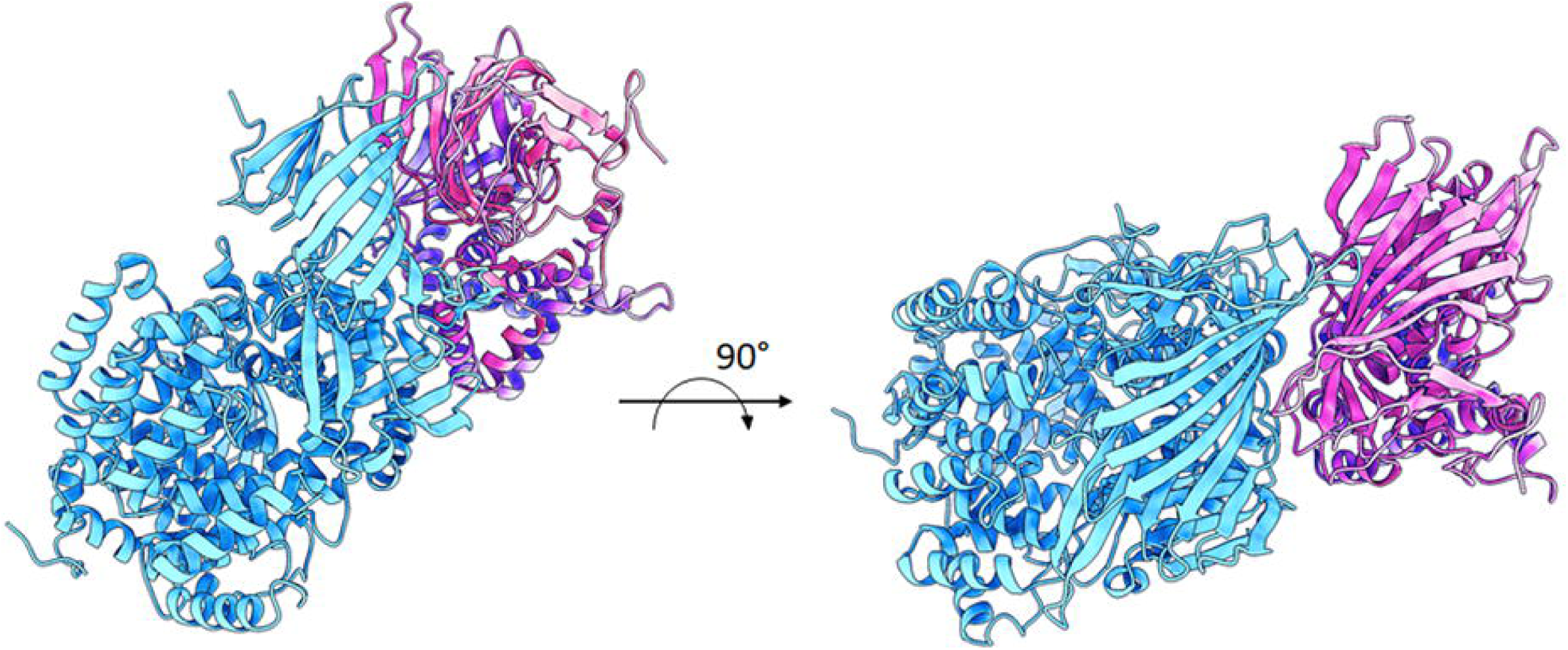
Proposed model of interaction between ERAP2 55kDa fragment and IRAP. ERAP2 (∼55kDa fragment; brown cartoons) and IRAP (cyan cartoons) are predicted to interact in a pseudo-symmetric heterodimer, with the N-terminal region of ERAP2 (residues 50-170) making extensive contacts with the corresponding region of IRAP.

### ERAP2 “short” derives from longer precursors

An open question is from which precursor the ∼55kDa fragment is generated. Indeed, while in individuals typed as “A” at rs2248374 as well in the case of the U937 cells, it is likely to derive from the full-length protein (Figures 1 and 4), in the case of the “G” haplotype we could not detect any precursor in macrophages. A possible explanation is that in macrophages, part of the ERAP2 mRNA copies can escape NMD and the naturally encoded truncated protein which normally undergoes proteolysis, can be stabilized, possibly by the binding to IRAP, when an acidic microenvironment allows the generation of the ∼55kDa fragment. To explore this hypothesis, two constructs were generated (Figure S1): one carrying the natural “G” mRNA which encodes for the specific sequence that stops at amino acid 532 and carries the sequence VRIKRVTE at C-terminus before the stop codon and the other carrying the alternative “A” mRNA but interrupted at the same length as the “G” haplotype and therefore differing at the C-terminus for the last amino acids (LAFLGENAEVK) (Figure 6). Following transfection into HEK293T cells, both proteins were expressed. However, the results show that, at low pH, the “short” ERAP2 is obtained only from the construct carrying the “VRIKRVTE” motif encoded by the “G” haplotype (Fig.6A). Modelling of the two truncated proteins support our findings: in the case of the “A” haplotype (LAFLGENAEVK) both the ion bridge between Asp517-Lys518 and the disulphide bridge between Cys503 and Cys514 are conserved whereas in the case of the “G” haplotype carrying at C-terminus the sequence “VRIKRVTE”, the disulphide bridge is lost and the conformation of the Asp517-Pro518 motif is kept away from Lys518, bringing the carboxyl side chain of Asp517 and the backbone amide on the neighbouring Pro518 into proximity, and locking the local moiety into an advantageous conformation for the subsequent hydrolysis at low pH values (Figure 6B).

**Figure 6.**
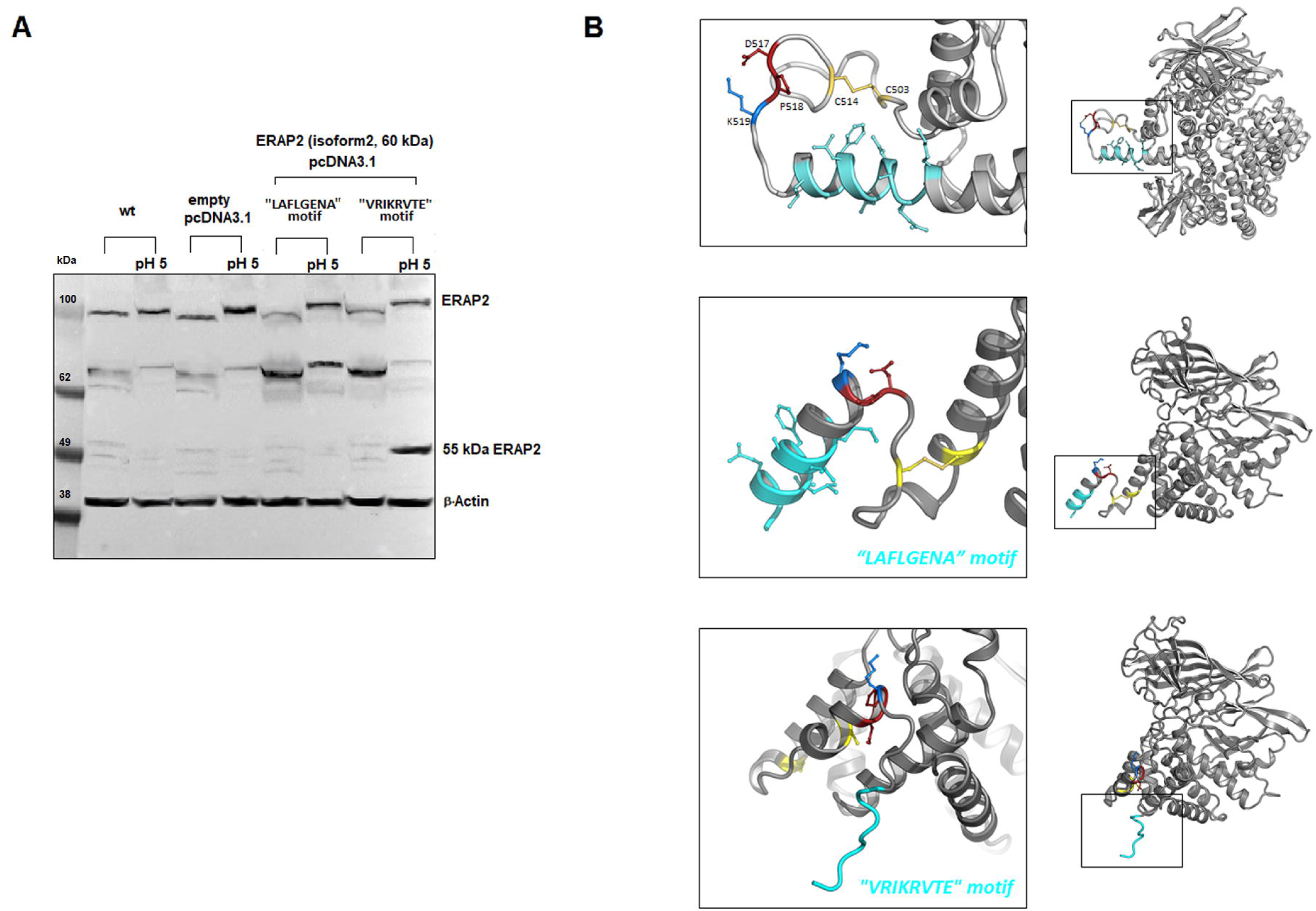
Expression and proposed model of the truncated proteins. **(A)** Western blot of the truncated proteins transfected into Hek293T cells. Plasmids containing the sequence reported in Fig. S1 corresponding to ERAP2/iso2 were transfected into Hek293T cells. The first 4 lanes are controls (Hek293T cells and Hek293T cells transfected with the vector pcDNA3.1. Lanes 5 and 6 report the Hek293T transfected with the plasmid expressing the 534 aa of the haplotype A (A at rs2248374) not treated or acid-treated lysate. Lanes 7 and 8 report the Hek293T transfected with the plasmid expressing the 532 aa corresponding to the haplotype G (G at rs2248374) not treated or acid-treated lysate. To note: when treated at pH5 before running, the samples run slightly higher than the untreated counterpart. **(B)** Modelling of the two truncated proteins. Rosetta 3.13 Linux version was used to predict the conformation of the two “short” ERAP2 carrying either the LAFLGENA or the VRIKRVTE motif. The initial atomic coordinates were taken from the X-ray crystal structure of ERAP2 (Protein Data Bank ID: 5AB0). The Python protocol described in Yang et al. (32), which makes use of the Rosetta 3.13 neural network to generate inter-residue distance and orientation constraints for a sequence of unknown structure, was used. The Rosetta minimizer was then used to find the backbone conformation consistent with the constraints.

We concluded that the specific C-terminus of the “G” isoform was responsible, in specific circumstances (i.e. acidic pH), and in some cells (i.e. macrophages or cancer cells), of the generation of the ∼55kDa fragment.

## DISCUSSION

We report here the existence of a short form of ERAP2 that binds IRAP and is secreted by the M2-type macrophages. Remarkably, this “short” ERAP2 is generated by an autocatalytic process that occurs in an acidic microenvironment. A similar event has been reported for MUC2 protein with which ERAP2 shares some features. Indeed, ERAP2 displays a unique disordered loop containing two cysteines (Cys503 and Cys514) and a close downstream Asp517-Pro518 specific sequence. The reduction of the disulfide bridge could expose and allow the hydrolysis of the Asp-Pro bond as previously described for the MUC2 in the late secretory pathway (40). It is known that IRAP lies in the endosomes whose acidic microenvironment can eventually determine the generation of the ERAP2 “short” fragment. We do not know, however, where exactly the encounter between ERAP2 and IRAP occurs: i.e. whether ERAP2 reaches the endosomes along its way to the cell membrane or whether it is endocytosed once bound to IRAP. Whatever the case, it is shown here that the two molecules are co-expressed in the endosomes suggesting that it is their acidic microenvironment that allows the catalytic process leading to the “short” ERAP2. In addition, the immunoprecipitation experiments, supported by the modelling, unambiguously demonstrate the existence of a direct, strong binding between the two molecules. It is interesting that the same phenomenon was observed in some cancer cell lines expressing the “short” ERAP2 although the secretion appears to be specific for the MDMs. Most remarkable however is the observation that, differently from the full-length protein, the “short” ERAP2 is expressed in macrophages independently from the polymorphism at rs2248374 that discriminates the two possible haplotypes. While this is conceivable in the haplotype expressing the full length ERAP2 (allele A at rs2248374), we could not observe any precursor to which the ∼55kDa fragment could be ascribed in the alternative haplotype. It is possible and not unlikely that in macrophages part of the mRNA copies escape NMD and the translated shorter form is stabilized only when the microenvironment conditions allow the formation of the ∼55KDa fragment as our data suggest. It must be noted that previous reports (41,42) have shown the existence of smaller ERAP2 isoform (isoform 3) of similar MW in macrophages carrying a G at rs2248374. The starting codon of the ERAP2/Iso3 however, maps after the terminal codon of ERAP2/Iso2, the latter corresponding to the sequence of the constructs expressed in Figure 6 and recognized by the anti-ERAP2 monoclonal antibody. Moreover, in the context of a previous work, we have sequenced the RNA from PMA-treated U937 cells (43) as well as from two monocytes/macrophages series (unpublished). No RNA corresponding to the ∼55kDa ERAP2 described here was found while the ERAP2 Iso/3, although listed, was undetectable in the U937 cells and barely detectable in monocytes and in the M1 but not in the M2 macrophages. These observations, together with the fact that the “short” ERAP2 is obtained by the acid-treatment of the cell lysates, strongly supports its catalytic origin.

Of note, our results also show that the process is distinctive of the differentiated myeloid cells within the PBMCs pointing to a specific role in these cells. It is known that IRAP binds with high affinity AT4, a biologically active fragment of Angiotensin II. IRAP is also named oxytocinase since it was found to regulate the level of circulating oxytocin during the later stages of human pregnancy and, interestingly, is also highly expressed in brain regions associated with cognition being recognized as a potential target for the treatment of cognitive disorders (44). The observation reported here that it can bind ERAP2 adds a novel piece to the intriguing puzzle regulating the IRAP multifaceted functions, including inflammation. In particular, it has been shown that ACE2 (Angiotensin I-converting enzyme-2) is the receptor for SARS-COV-2 (45) and it has been postulated that its reduction in the cell membrane due to the endocytic pathway associated with SARS-CoV-2 binding, can play a role in altering the inflammatory balance (16). ERAP2 “short” fragment secreted by type 2 macrophages, by binding IRAP is likely to interfere with the RAS (Renin-Angiotensin-System) by modulating its anti-inflammatory arm. In this context, a polymorphism in the ERAP2 gene has been found associated with death due to SARS-CoV-2 (46). The possible role of this molecule in the SARS-CoV-2 infection is definitively worth investigating further.

In addition to this and most interestingly, our results offer a new sight on the peculiar evolutionary story of ERAP2. Indeed, this gene is absent in many species and, so far, believed to be redundant in humans since at least a quarter of subjects does not express ERAP2. We have shown here that, even in macrophages not expressing the full length ERAP2, this is present as a fragment in specialized cells, playing a role that, although in need of further investigation, appears to be unrelated to the conventional antigen presentation. It is also conceivable that this role is the reason why the *ERAP2* gene has been retained in humans under a balanced selection so that this fragment can be expressed by everyone independently from the haplotype, even though ERAP2 full length remains dispensable. Indeed, most of the reported *ERAP2*-disease associations are with the haplotype expressing the full-length protein whose presence might therefore even be detrimental. Interestingly, we observed that monocytes stimulated for several days with M-CSF as well as PMA-treated U937 cells, tend to acquire a M2 phenotype as shown by the expression of the marker MS4A4A, and therefore secrete the “short” ERAP2. This allows to speculate that ERAP2, through the binding with IRAP, can be secreted by aging macrophages and possibly, through IRAP, could keep at bay the inflammation associated with senescence (47). A further interesting observation is that this feature, i.e. the expression of the ∼55kDa ERAP2 and its binding to IRAP, is shared with some cancer cell lines. It is well known that transformed cells prefer a metabolic pathway leading to an acidic microenvironment which can promote the generation of the ∼55kDa fragment. However, we could not observe this fragment in the surnatant suggesting that the secretion might be specific for the type 2 macrophages.

In conclusion, the observations reported here, although not conclusive about the exact mechanisms underpinning the generation and the activity of this “short” ERAP2, are pivotal in several contexts thus opening new fields of investigation.

## AUTHOR CONTRIBUTIONS

FP, BM and RS designed and performed experiments; analyzed data and drafted the manuscript. FP produced and analyzed the genetic data and flow cytometry assay; BM and SC did the confocal analysis; FP, BM, SD, EV and MF processed the samples, developed and performed western blots and analysis. AP did the bioinformatic models. All authors critically reviewed the manuscript and approved the submitted version.

## ACKNOWLEDGEMENTS

The authors wish to thank Valentina Tedeschi, Maria Teresa Fiorillo and all the members of the laboratory for suggestions and are grateful to Federica Lucantoni for the excellent technical assistance. This work was supported by Ceschina Foundation, Switzerland and Sapienza University of Rome, Italy.

## References

1. Tsujimoto M & Hattori A. The oxytocinase subfamily of M1 aminopeptidases. Biochim Biophys Acta (2005) 1751:9–18. doi: 10.1016/j.bbapap.2004.09.011

2. Paladini F, Fiorillo MT, Tedeschi V, Mattorre B & Sorrentino R. The Multifaceted Nature of Aminopeptidases ERAP1, ERAP2, and LNPEP: From Evolution to Disease. Front Immunol (2020) 11:1576. doi: 10.3389/fimmu.2020.01576. eCollection 2020.

3. Weimershaus M, Evnouchidou I, Saveanu L & van Endert P. Peptidases trimming MHC class I ligands. Curr Opin Immunol (2013) 25:90−96. doi: 10.1016/j.coi.2012.10.001.

4. Saveanu L, Carroll O, Weimershaus M, Guermonprez P, First E, Lindo V, et al. IRAP identifes an endosomal compartment required for MHC class I cross-presentation. Science (2009) 325:213–217. doi: 10.1126/science.1172845.

5. Weimershaus M, Mauvais FX, Evnouchidou I, Lawand M, Saveanu L & van Endert P. IRAP Endosomes Control Phagosomal Maturation in Dendritic Cells. Front Cell Dev Biol (2020) 8:585713. doi: 10.3389/fcell.2020.585713. eCollection 2020.

6. Vear A, Gaspari T, Thompson P & Chai SY. Is There an Interplay Between the Functional Domains of IRAP? Front Cell Dev Biol (2020) 8:585237. doi: 10.3389/fcell.2020.585237.

7. Descamps D, Evnouchidou I, Caillens V, Drajac C, Riffault S, van Endert P et al. The Role of Insulin Regulated Aminopeptidase in Endocytic Trafficking and Receptor Signaling in Immune Cells. Front Mol Biosci (2020) 7:583556. doi: 10.3389/fmolb.2020.583556. eCollection 2020.

8. Albiston AL, Morton CJ, Ng HL, Pham V, Yeatman HR et al. Identification and characterization of a new cognitive enhancer based on inhibition of insulin-regulated aminopeptidase. FASEB J (2008) 22:4209–17. doi: 10.1096/fj.08-112227.

9. Vargas F, Wangesteen R, Rodríguez-Gómez I & García-Estañ J. Aminopeptidases in Cardiovascular and Renal Function. Role as Predictive Renal Injury Biomarkers. Int J Mol Sci (2020) 21:5615. doi: 10.3390/ijms21165615.

10. Sharip A & Kunz J. Understanding the Pathogenesis of Spondyloarthritis. Biomolecules (2020) 10:1461. doi: 10.3390/biom10101461.

11. Babaie F, Hosseinzadeh R, Ebrazeh M, Seyfizadeh N, Aslani S, Salimi S et al. The roles of ERAP1 and ERAP2 in autoimmunity and cancer immunity: New insights and perspective. Mol Immunol (2020) 121:7–19. doi: 10.1016/j.molimm.2020.02.020.

12. Tedeschi V, Paldino G, Paladini F, Mattorre B, Tuosto L, Sorrentino R et al. The Impact of the ‘Mis-Peptidome’ on HLA Class I-Mediated Diseases: Contribution of ERAP1 and ERAP2 and Effects on the Immune Response. Int J Mol Sci (2020) 21:9608. doi: 10.3390/ijms21249608.

13. Zhen Q, Yang Z, Wang W, Li B, Bai M, Wu J et al. Genetic Study on Small Insertions and Deletions in Psoriasis Reveals a Role in Complex Human Diseases. J Invest Dermatol (2019) 139:2302-2312.e14. doi: 10.1016/j.jid.2019.03.1157.

14. Li DT, Habtemichael EN & Bogan JS. Vasopressin inactivation: Role of insulin-regulated aminopeptidase. Vitam Horm (2020) 113:101–128. doi: 10.1016/bs.vh.2019.08.017.

15. Compagnone M, Cifaldi L & Fruci D. Regulation of ERAP1 and ERAP2 genes and their disfunction in human cancer. Hum Immunol (2019) 80:318–324. doi: 10.1016/j.humimm.2019.02.014.

16. D’Amico S, Tempora P, Lucarini V, Melaiu O, Gaspari S, Algeri M et al. ERAP1 and ERAP2 Enzymes: A Protective Shield for RAS against COVID-19? Int J Mol Sci (2021) 22:1705. doi: 10.3390/ijms22041705.

17. Lee ED. Endoplasmic Reticulum Aminopeptidase 2, a common immunological link to adverse pregnancy outcomes and cancer clearance? Placenta (2017) 56:40–43. doi: 10.1016/j.placenta.2017.03.012.

18. Soltani S & Nasiri M. Association of ERAP2 gene variants with risk of pre-eclampsia among Iranian women. Int J Gynaecol Obstet (2019) 145:337–42. doi: 10.1002/ijgo.12816.

19. McGonagle D, Aydin SZ, Gül A, Mahr A, § Direskeneli H. MHC-I-opathy-unified concept for spondyloarthritis and Behçet disease. Nat Rev Rheumatol (2015) 11:731–740. doi: 10.1038/nrrheum.2015.14

20. Kenna TJ, Robinson PC & Haroon N. Endoplasmic reticulum aminopeptidases in the pathogenesis of ankylosing spondylitis. Rheumatology (2015) 54:1549–56. doi: 10.1093/rheumatology/kev218.

21. Andrés AM, Dennis MY, Kretzschmar WW, Cannons JL, Lee-Lin SQ et al. Balancing selection maintains a form of ERAP2 that undergoes nonsense-mediated decay and affects antigen presentation. PLoS Genet (2010) 6:e1001157. doi: 10.1371/journal.pgen.1001157.

22. Goto Y, Ogawa K, Hattori A & Tsujimoto M. Secretion of endoplasmic reticulum aminopeptidase 1 is involved in the activation of macrophages induced by lipopolysaccharide and interferon-gamma. J Biol Chem (2011) 286:21906–21914. doi: 10.1074/jbc.M111.239111.

23. Ofner LD & Hooper NM. Ectodomain shedding of cystinyl aminopeptidase from human placental membranes. Placenta (2002) 1:65–70. doi: 10.1053/plac.2001.0751.

24. Saulle I, Ibba SV, Torretta E, Vittori C, Fenizia C, Piancone F, et al. Endoplasmic Reticulum Associated Aminopeptidase 2 (ERAP2) Is Released in the Secretom,e of Activated MDMs and Reduces in vitro HIV-1 Infection. Front Immunol (2019) 10:1648. doi: 10.3389/fimmu.2019.01648. eCollection 2019.

25. Paladini F, Fiorillo MT, Vitulano C, Tedeschi V, Piga M, Cauli A, et al. An allelic variant in the intergenic region between ERAP1 and ERAP2 correlates with an inverse expression of the two genes. Sci Rep (2018) 8, 10398 28799–8.

26. Fic E, Kedracka-Krok S, Jankowska U, Pirog A & Dziedzicka-Wasylewska M. Comparison of protein precipitation methods for various rat brain structures prior to proteomic analysis. Electrophoresis (2010) 21:3573–3579. doi: 10.1002/elps.201000197.

27. Schneider C, Rasband W & Eliceiri K. NIH Image to ImageJ: 25 years of image analysis. Nat Methods (2012) 9:671–675. doi: 10.1038/nmeth.2089.

28. Zinchuk V, Zinchuk O§ Okada T. Quantitative Colocalization Analysis of Multicolor Confocal Immunofluorescence Microscopy Images: Pushing Pixels to Explore Biological Phenomena. Acta Histochem. Cytochem. (2007) 40:101–111

29. Kochan G, Krojer T, Harvey D, Fischer R, Chen L, Vollmar M et al. Crystal structures of the endoplasmic reticulum aminopeptidase-1 (ERAP1) reveal the molecular basis for N-terminal peptide trimming. Proc Natl Acad Sci USA (2011) 108:7745–50. doi: 10.1073/pnas.1101262108.

30. Mpakali A, Giastas P, Mathioudakis N, Mavridis IM, Saridakis E et al. Structural Basis for Antigenic Peptide Recognition and Processing by Endoplasmic Reticulum (ER) Aminopeptidase 2. J Biol Chem (2015) 290:26021–26032. doi: 10.1074/jbc.M115.685909.

31. Hermans SJ, Ascher DB, Hancock NC, Holien JK, Michell BJ, Chai SY, et al. Crystal structure of human insulin-regulated aminopeptidase with specificity for cyclic peptides. Protein Sci (2015). 24:190–199. doi: 10.1002/pro.2604.

32. de Vries SJ, van Dijk M & Bonvin AM. The HADDOCK web server for data-driven biomolecular docking. Nat Protoc (2010) 5:883–897. doi: 10.1038/nprot.2010.32.

33. Vangone A & Bonvin AMJJ. Contact-based prediction of binding affinity in protein-protein complexes. eLife (2015) 4:e07454. doi: 10.21769/BioProtoc.2124.

34. Xue L, Rodrigues J, Kastritis P, Bonvin AMJJ & Vangone A. PRODIGY: a web-server for predicting the binding affinity in protein-protein complexes. Bioinformatics (2016) 32:3676–3678. doi:10.1093/bioinformatics/btw514.

35. Yang J, Anishchenko I, Park H, Peng Z, Ovchinnikov S & Baker D. Improved protein structure prediction using predicted interresidue orientations. PNAS (2020) 117:1496–1503; doi: 10.1073/pnas.1914677117.

36. Mattiola I, Tomay F, De Pizzol M, Silva-Gomes R, Savino B, Gulic T et al. The macrophage tetraspan MS4A4A enhances dectin-1-dependent NK cell-mediated resistance to metastasis. Nat Immunol (2019) 8:1012–1022. doi: 10.1038/s41590-019-0417-y.

37. Fruci D, Ferracuti S, Limongi MZ, Cunsolo V, Giorda E, Fraioli R et al. Expression of Endoplasmic Reticulum Aminopeptidases in EBV-B Cell Lines from Healthy Donors and in Leukemia/Lymphoma, Carcinoma, and Melanoma Cell Lines. J Immunol (2006) 176:4869–4879. doi: 10.4049/jimmunol.176.8.4869.

38. Sanyal R, Polyak MJ, Zuccolo J, Puri M, Deng L, Roberts L et al. MS4A4A: a novel cell surface marker for M2 macrophages and plasma cells. Immunol Cell Biol (2017) 95:611–619. doi: 10.1038/icb.2017.18.

39. Papakyriakou A & Stratikos E. The Role of Conformational Dynamics in Antigen Trimming by Intracellular Aminopeptidases. Front Immunol (2017) 8:946. doi: 10.3389/fimmu.2017.00946.

40. Lidell ME, Johansson MEV & Hansson GC. An autocatalytic cleavage in the C terminus of the human MUC2 mucin occurs at the low pH of the late secretory pathway. J. Biol Chem (2003) 278:13944–13951. doi: 10.1074/jbc.M210069200.

41. Saulle I, Vanetti C, Goglia S, Vicentini C, Tombetti E, Garziano M et al. A New ERAP2/Iso3 Isoform Expression Is Triggered by Different Microbial Stimuli in Human Cells. Could It Play a Role in the Modulation of SARS-CoV-2 Infection? Cells (2020) 9:1951. doi: 10.3390/cells9091951.

42. Ye CJ, Chen J, Villani AC, Gate RE, Subramaniam M, Bhangale T et al. Genetic analysis of isoform usage in the human anti-viral response reveals influenza-specific regulation of ERAP2 transcripts under balancing selection. Genome Res (2018) 28:1812–1825. doi: 10.1101/gr.240390.118.

43. Rossetti C, Picardi E, Ye M, Camilli G, D’Erchia AM, Cucina L. et al. RNA editing signature during myeloid leukemia cell differentiation. Leukemia (2017) 12:2824–2832. doi: 10.1038/leu.2017.134.

44. Georgiadis D, Ziotopoulou A, Kaloumenou E, Lelis A & Papasava A. The Discovery of Insulin-Regulated Aminopeptidase (IRAP) Inhibitors: A Literature Review. Front Pharmacol (2020) 11:585838. doi: 10.3389/fphar.2020.585838. eCollection 2020.

45. Yang J, Petitjean SJL, Koehler M, Zhang Q, Dumitru AC, Chen W et al. Molecular interaction and inhibition of SARS-CoV-2 binding to the ACE2 receptor. Nat Commun (2020) 11:4541. doi: 10.1038/s41467-020-18319-6.

46. Lu C, Gam R, Pandurangan AP & Goug J. Genetic risk factors for death with SARS-CoV-2 from the UK Biobank. medRxiv (2020). doi: 10.1101/2020.07.01.20144592.

47. Barbé-Tuana F, Funchal G, Schmitz CRR, Maurmann RM & Baue ME. The interplay between immunosenescence and age-related diseases. Semin Immunopathol (2020) 42:545–557. doi: 10.1007/s00281-020-00806-z.

