## supplementary data for "A short ERAP2 that binds IRAP is expressed in macrophages independently from gene variation"

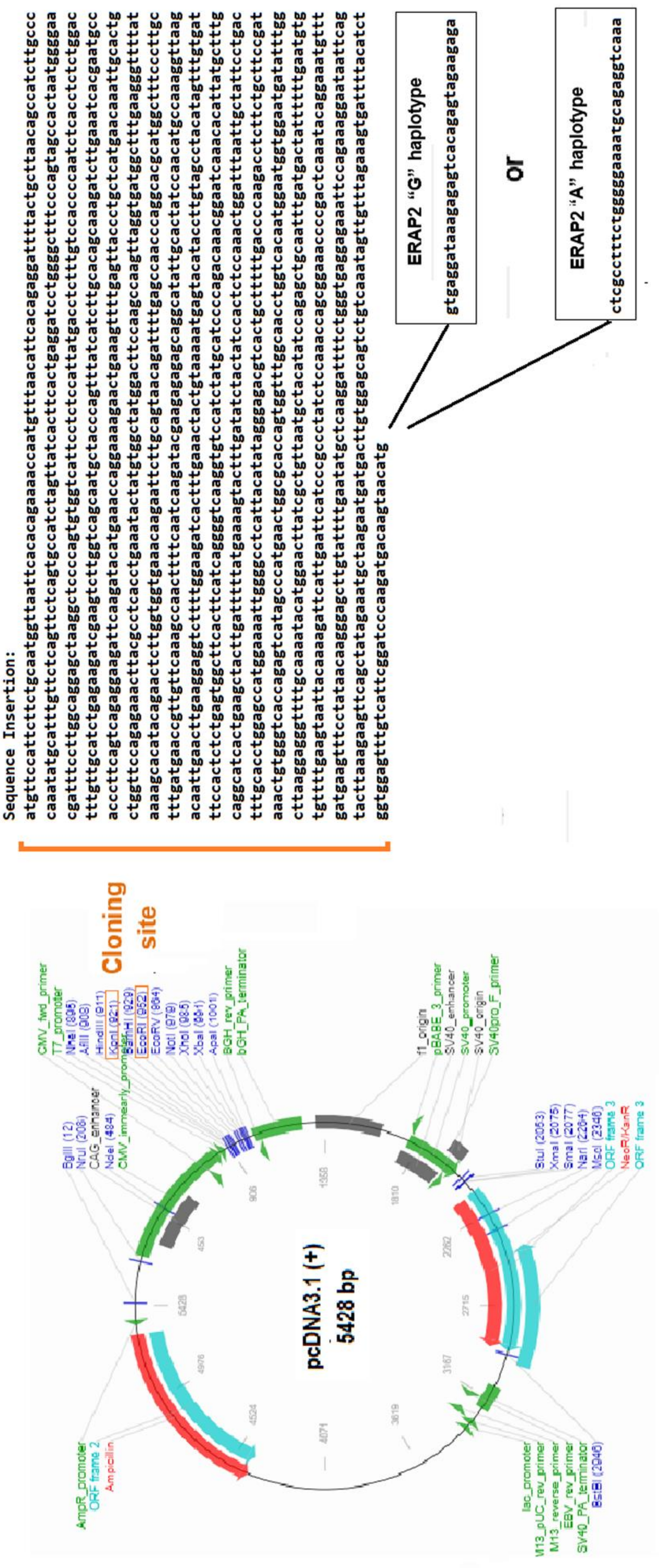


**Figure S1:** Detailed sequences of the constructs carrying the ERAP2 derived from the "G" or the "A" haplotype encoding the first 532 or 534 aminoacids respectively.

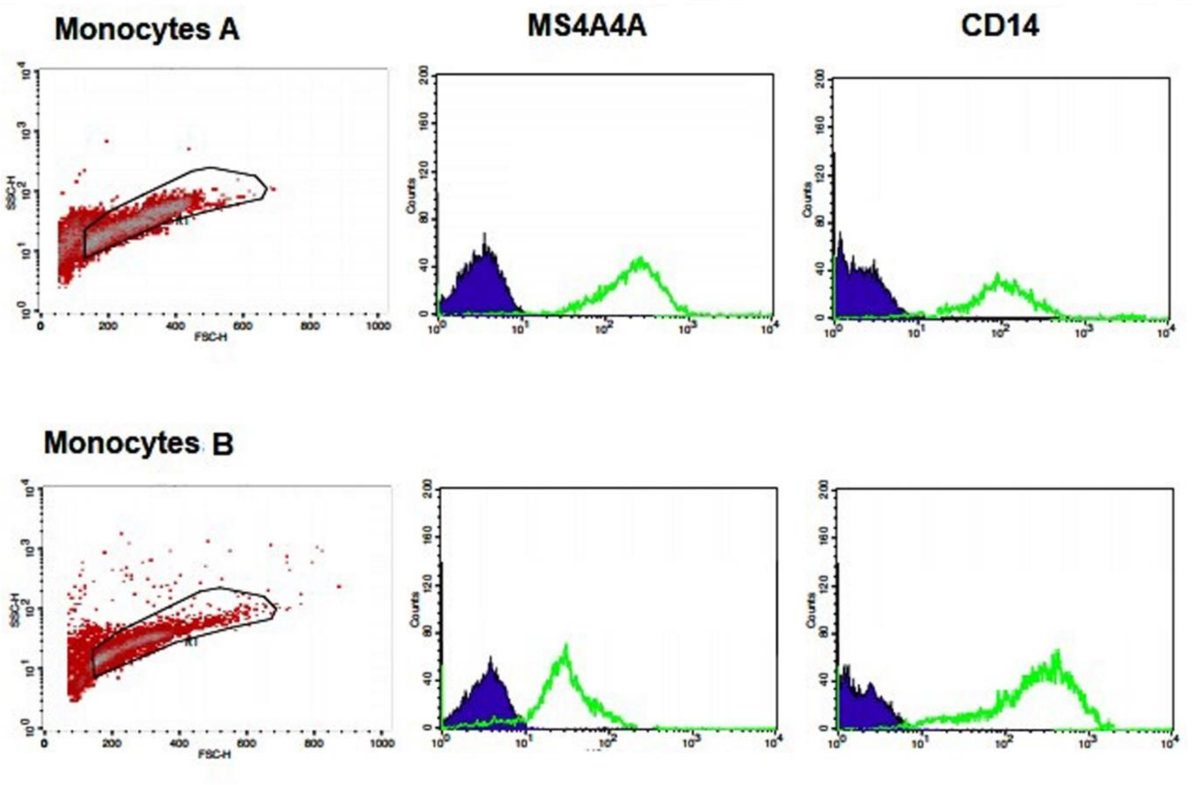


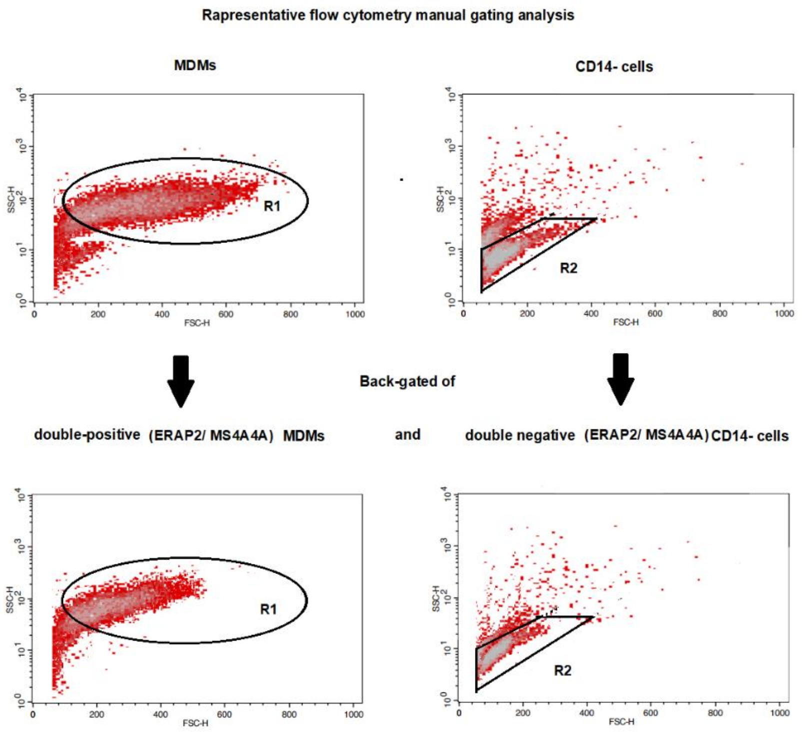


**Figure S2:** Representative cytofluorimetric analysis of the expression of MS4A4A in monocytes genotyped as A/A (A) or G/G (B) at rs2248374.

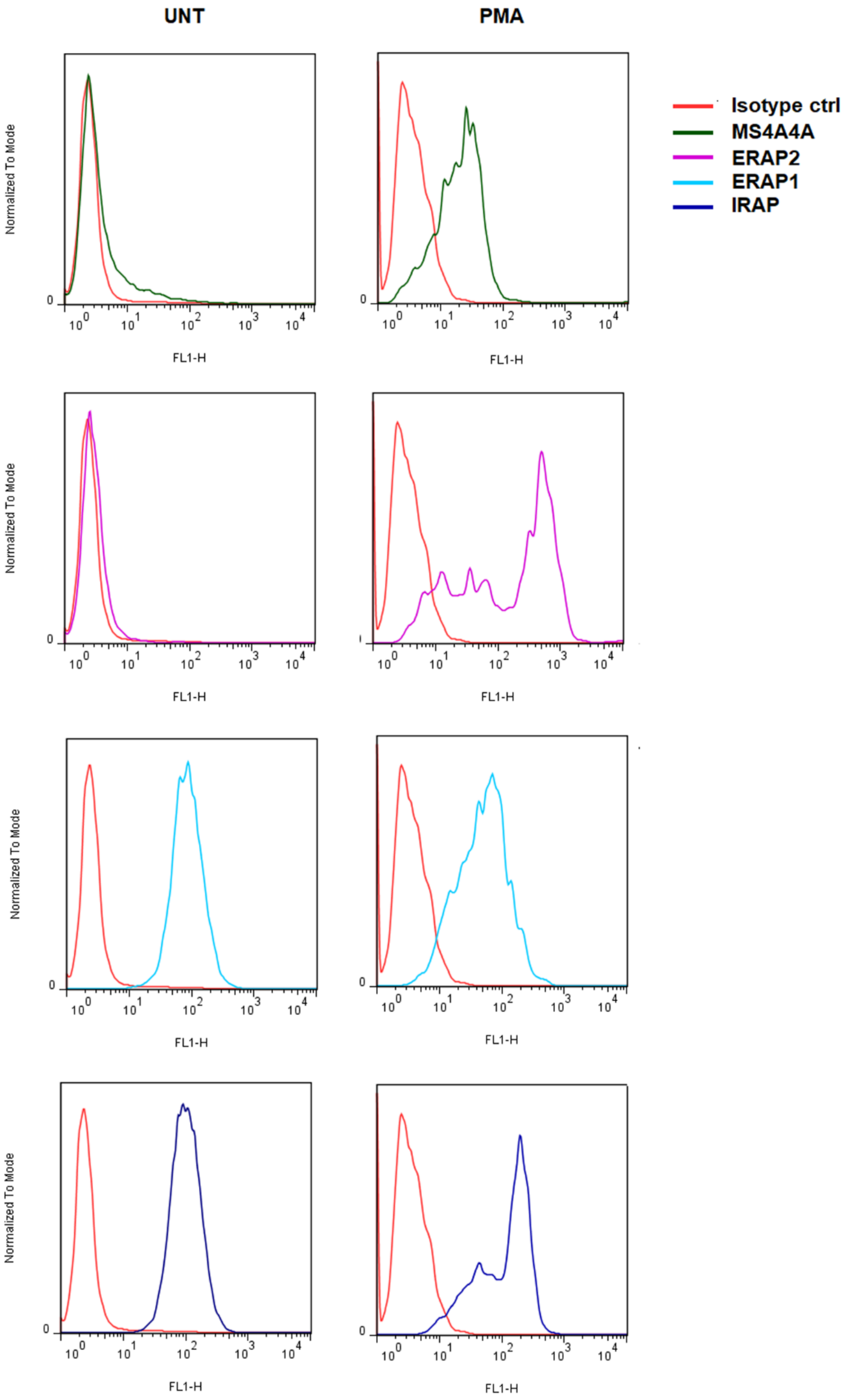


**Figure S3:** Cytofluorimetric analysis of the expression of MS4A4A, ERAP2, ERAP1 and IRAP in untreated (left panels) and PMA-treated (right panels) U937 cells.

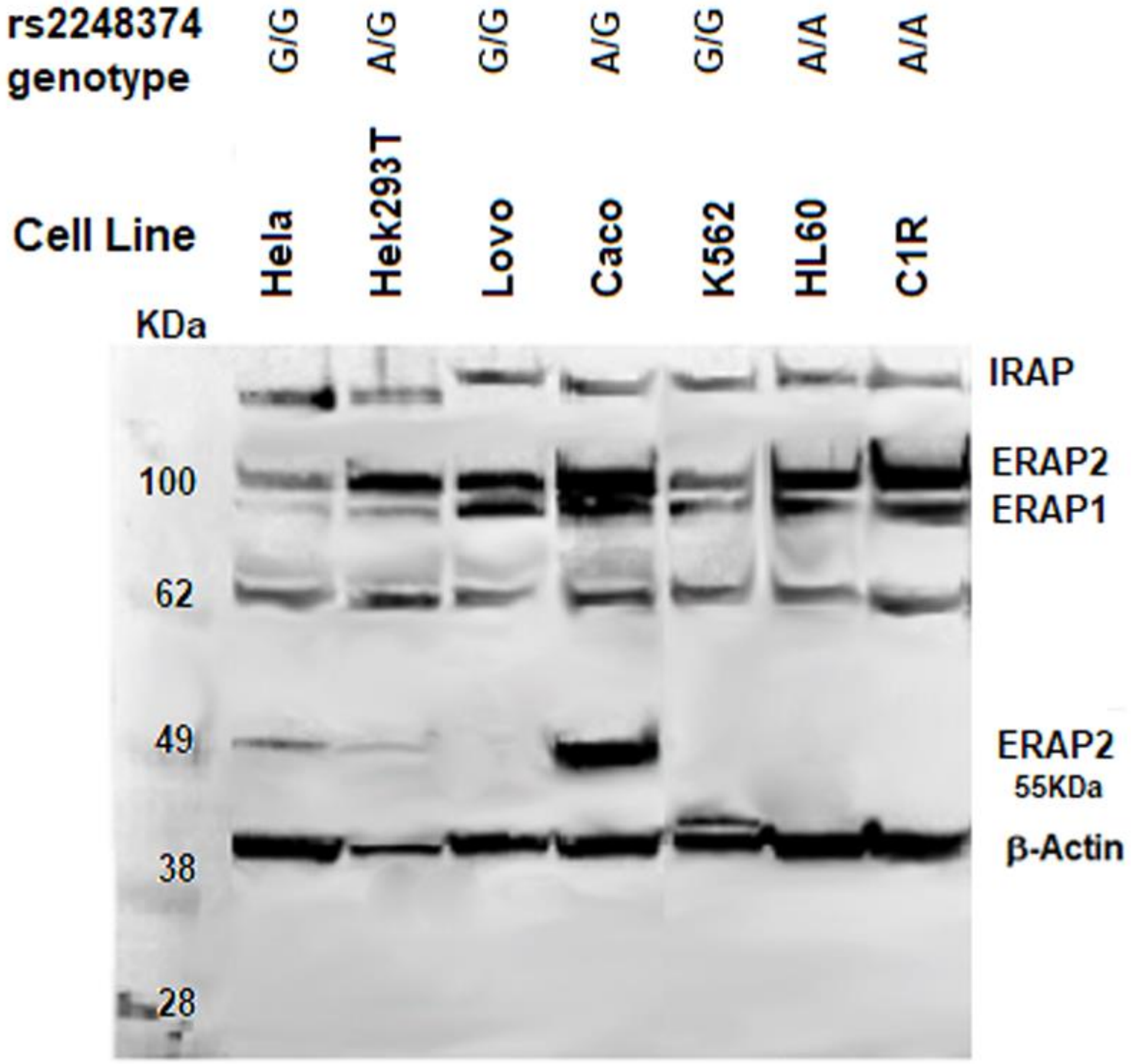


**Figure S4:** ERAP2, ERAP1 and IRAP expression in cancer cell lines. The respective rs2248374 genotypes are indicated at the top of the image.
